## Supplementary Figure 1-5, Supplementary Table 1 for "Direct visualization of human myosin II force generation using DNA origami-based thick filaments"

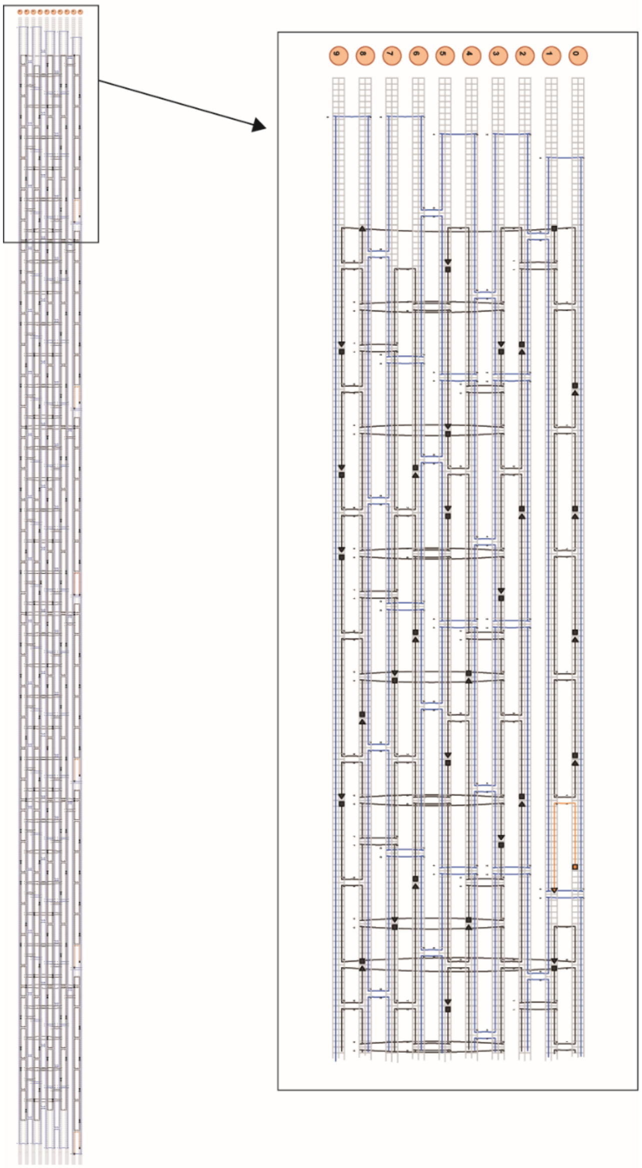

**Supplementary Figure 1. Design of the DNA origami rod for the engineered thick filament** The scheme was produced by caDNAno2 software (https://cadnano.org/). Scaffold, core staples, handles for linking with myosin and the actin binding domain of α-actinin are shown in blue, black, red and orange, respectively. The boxed area is expanded on the right.

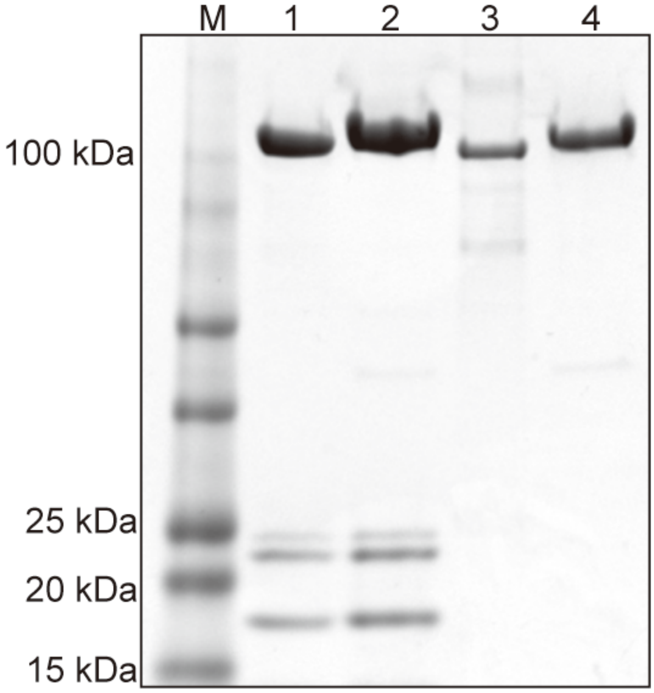


**Supplementary Figure 2. SDS-PAGE gel of human myosin IIa S1 and lever-arm-less S1** Myosin S1 showed light chains (15-25 kDa), lever-arm-less S1 did not. A band shift for heavy chains (~100 kDa) confirmed the labeling of oligonucleotides. M, marker; lane 1, myosin IIa S1; lane 2, myosin IIa S1 labeled with a 21 mer oligonucleotide; lane 3, lever-arm-less S1; lane 4, lever-arm-less S1 labeled with a 21 mer oligonucleotide.

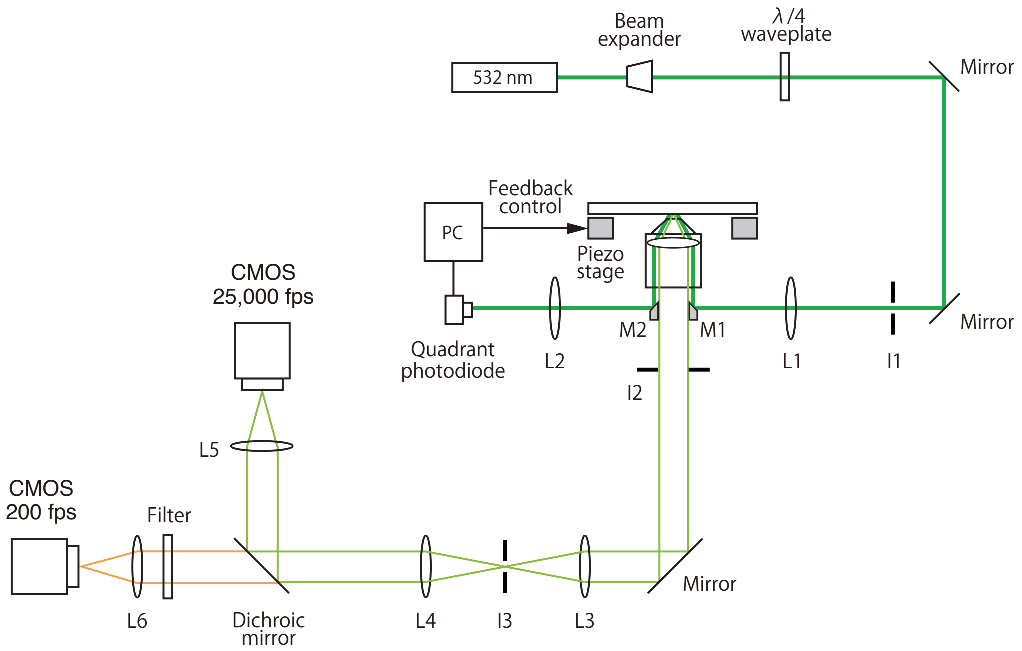


**Supplementary Figure 3. A schematic of the TIRF setup** A 532 nm laser was collimated and expanded by a beam expander consisting of two lenses. A λ/4 waveplate was used to achieve circular polarized illumination. An iris (I1) was used to reduce the illumination area. The laser was focused onto the back focal plane of the objective by a lens (L1, 250 mm focal length) and micromirror (M1). The laser reflected on a glass surface immediately below the sample was directed into a quadrant photodiode by a micromirror (M2) and lens (L2) to perform feedback control of the piezo stage. An iris (I2) was used to attenuate scattering light. Scattering and fluorescent light were collimated by two lenses (L3 and L4), separated by a dichroic mirror and finally imaged onto CMOS cameras by imaging lenses (L5 and L6). An iris (I3) was used to define the field of view. A filter was used to increase the signal-to-noise ratio of the fluorescence image. See Methods for details.

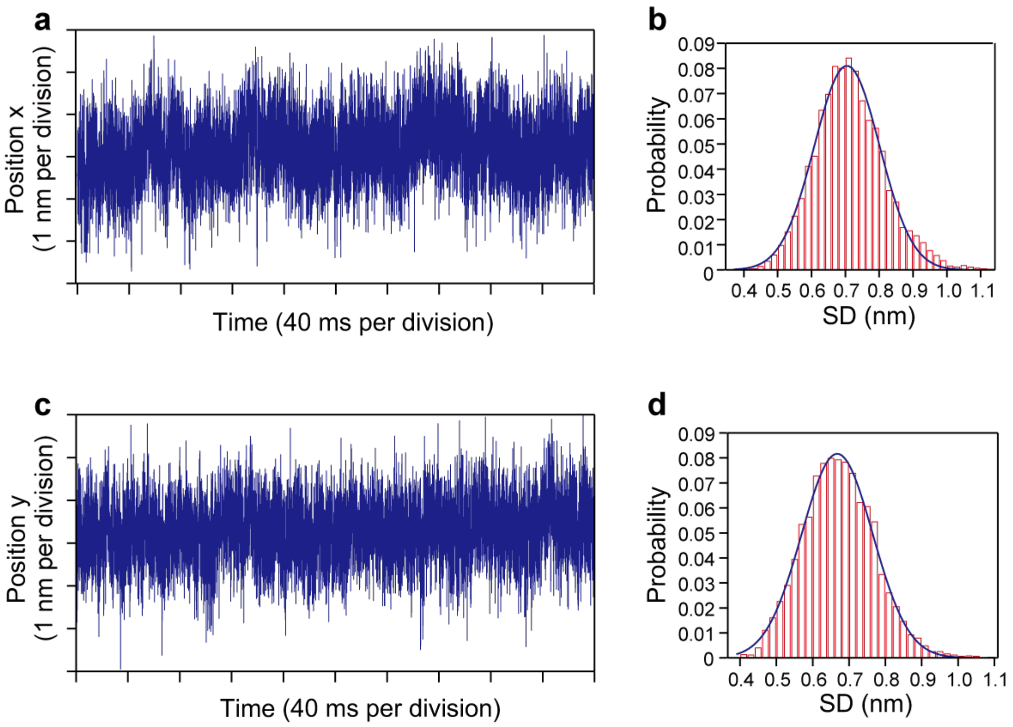


**Supplementary Figure 4. A representative trace of an immobile GNP on a glass surface** An immobile GNP on a glass surface was imaged at 25,000 fps for 400 ms and localized. The drift of the stage was corrected by tracing a few immobile GNPs in the field of view, averaging the traces and subtracting from the raw trace. The GNP was immobilized on a glass surface via electrostatic interactions. **a**, **c** GNP trajectories in the x-direction (**a**) and y-direction (**c**). **b**, **d** Histograms of the standard deviations of the trajectories (**b**, x-direction; **d**, y-direction). The standard deviations were calculated every 25 points. The peaks are (x-direction) 0.70 ± 0.10 nm and (y-direction) 0.67 ± 0.10 nm (mean ± SD).

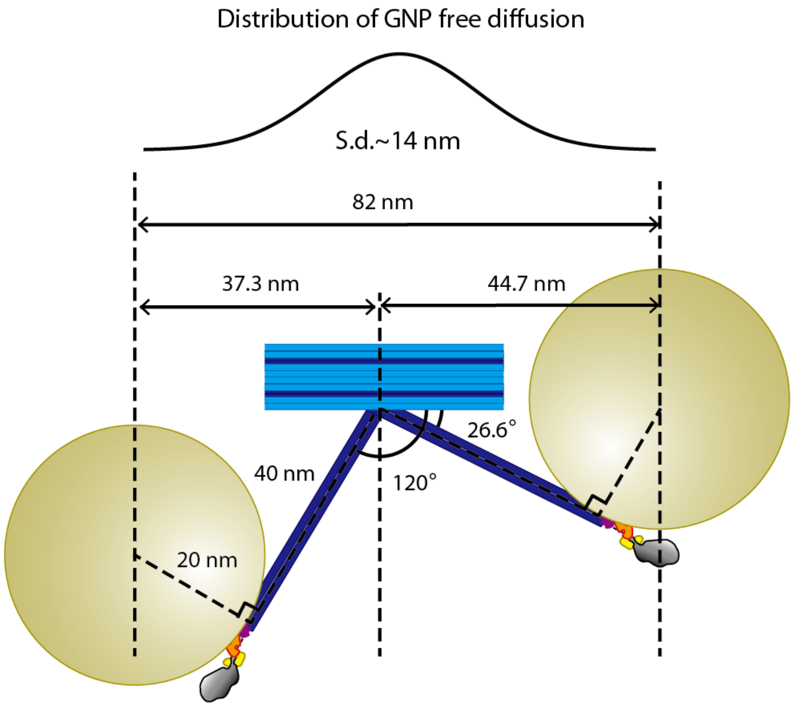


**Supplementary Figure 5. A schematic geometry of the engineered thick filament-myosin-GNP complex** The range of free diffusion of a GNP attached to the engineered thick filament was restricted by the structure of the engineered thick filament. The right side is the direction of the lever-arm swing. The S2-like linker, which is 40 nm long, can pivot from 0° to 120° against the backbone (Fig. 1e). Consequently, the center of the GNP can freely diffuse from 26.6° to 120° on the assumption that it is rigidly attached to the end of the S2-like linker via bifunctional DNA linkers (Fig. 3b), and S1 can flexibly rotate against the S2-like linker because of a nick in the DNA. The full range of GNP is 82 nm and, assuming a normal distribution of GNP free diffusion, the standard deviation is 14 nm. Our S2-like linker is approximately 3 times more flexible than native S2 (~20 nm peak-to-peak diffusion for native S2 tip[24] vs. ~60 nm peak-to-peak diffusion for our S2 tip).

**Supplementary Movie 1. High-speed AFM movie showing the two-step lever-arm swing of myosin II**

The dynamic process in 1 μM ATP was filmed at 400 ms $frame^{-1}$ (2.5 frames $s^{-1}$). Orientation changes in myosin II S1 were observed in a two-step manner. Image area, 66 × 96 $nm^2$ with 220 × 320 pixels (magnified 10 times for clarity by image processing).

**Supplementary Movie 2. High-speed AFM movie showing reversed lever-arm swing of myosin II**

The dynamic process at 1 μM caged ATP was filmed at 400 ms $frame^{-1}$ (2.5 frames $s^{-1}$). Image area, 66 × 81 $nm^2$ with 220 × 270 pixels (magnified 10 times for clarity by image processing).

**Supplementary Table 1. Oligonucleotides sequences for DNA rod**

This file contains the sequences of the core staples used to build the DNA rod for the synthetic thick filament and the sequences of the handle staples attached to myosin and the actin binding domain of α-actinin.

### Supplementary Table 1

Core staples to build the DNA rod for the synthetic thick filament.

| |
|---|
| ATGCCCCCTGCCTAGGTCAGACGATTGGCCTTGATACAGTTA |
| CAGTGCCCGTATAAATTCACAAACAAATAAATCCTTGAGTAA |
| CGGGGTCAGTGCCTCATTAAAGCCAGAATGGAAAGGTTTTAA |
| TGTACTGGTAATAACGCAGTCTCTGAATTTACCGTACAGGAG |
| AATAAACAGCCATAAATCAGATATAGAAGGCTTATGTTACAA |
| AGCCTAATTTGCCACCGGTATTCTAAGAACGCGAGTTTCCAG |
| CGCTAACGAGCGTCGCGTTTTAGCGAACCTCCCGATTACCAA |
| TTTTATCCTGAATCCTTGCGGGAGGTTTTGAAGCCGCTACAA |
| TTGCTTTGAATACCATCAATATAATCCTGATTGTTCGCCTGA |
| AACAATAACGGATTTGGATTATACTTCTGAATAATCGGGAGA |
| AGTACCTTTTACATGGAAGGGTTAGAACCTACCATCAGTAAC |
| GTCAGATGAATATAATCAAAATTATTTGCACGTAAGTTTAAC |
| CAGTGAGACGGGCACGAGCCGGAAGCATAAAGTGTTTTTCAC |
| CAGGGTGGTTTTTCAAAGCCTGGGGTGCCTAATGATGGGCGC |
| GGCGGTTTGCGTATGTGAGCTAACTCACATTAATTGGGGAGA |
| ATCGGCCAACGCGCGCGTTGCGCTCACTGCCCGCTTTAATGA |
| ACGGCCAGTGCCAATCCGGCACCGCTTCTGGTGCCTAAAACG |
| CAGTCACGACGTTGGGAAACCAGGCAAAGCGCCATGTTTTCC |
| TGGGTAACGCCAGGTCGCCATTCAGGCTGCGCAACATTAAGT |
| TGTGCTGCAAGGCGTGTTGGGAAGGGCGATCGGTGAGGGGGA |
| AACTAAAGTACGGTCCCGAAAGACTTCAAATATCGAATATGC |
| CTCAACATGTTTTACGTTTTAATTCGAGCTTCAAAGCTGTAG |
| TTGCTGAATATAATGCGAACCAGACCGGAAGCAAAAGCTTAA |
| TGCGGATGGCTTAGCTCCAACAGGTCAGGATTAGATCATTTT |
| AGGAGGTGACAGGCGGGGTTTTGCTCACAATCACCCCACCCTCATTTTC |
| TACAATAGCGCCACGAATAACATAAAGCGAGCATGTCTAAAGTTTTGTC |
| ATTGATGTCAGATAAAGAAAACAAAAGACAATTCGAATTTTTTCACGTT |
| TGCCACACAATTCACGAAAATCCTGTAGTTCTGCCGACAACAACCATCG |
| CATCCAGCGCACCCTTACCAGTCCCGAGGGATTCTGGCTACAGAGGCTT |
| TTATTAAAAAAGCTACATTTCGCAAACTATCGTCACTCATCTTTGACCC |
| CCGCCGCCAGCATTTGAGGCATTTCGGAACCTATTAGAGCCAGGTACCG |
| ATTACCAATCGATAGCAGCGGGGCGACAGTACCGT |
| CCCAAAAGAACTGGAGCAAACGTAGACGAGAGCAACCTGTAG |
| TAGGAATCATTACCGCAAGCATTATTTATCCCAATGTAACGATATAACC |
| AGGCATTGACAAAAGGTAAATTACTAGATCTGTAT |
| TTATATAACTATATAAGAACGCGAGACTCCTTAGAGAGTGAG |
| CATATTCCTGATTAGCAATTCAAGTTACAAAATCGGAATAATACGCTTC |

| |
|---|
| ACCTCAAGGCAAATCAACAGCAAAAATAGGAGCCT |
| ATTATTTACATTGGGGGACATTCTGGTTAACGTCAGAATTTC |
| TTGTTATCCGCTCACAACATAACAGCTGATTGCCCCGACAATAGAACAA |
| GGTGTGTTTACCTGCAGCCGTCGGGGTCTGAGGCT |
| CTCGTCGCTGGCAGGTGCCGGACTTGCCGTTGTGTACCCTCA |
| GGCCTCAGGAAGATCCAGCTTGCTTTCAGAGGTGGTAGCAACCCAATTT |
| GCAAATATTCGCATTAAATCACAGGTCAGTTTCCA |
| TTTAGAACCCTCATGGTAAAGATTCATTATTAGCACACCAAC |
| AGCGGATTGCATCAGAGGAAGGTCTGGAAGTTTCATAAAACATAAACAT |
| ACATAACACACTATCATAAGAATTATACGTACAAC |
| ACCACCGGTCCCTCAGAGCCGCCAGTACCAGGCGGCAGAACC |
| GCCAGTAATACTTGAGCCATTTGACCGTAATCAGTTTAAAGG |
| GCAATGCATTACCAGAAGGAAACAAATACATACATAAAGTAA |
| ATTTTGAAATCGAGAACAAGCAAAACAGGGAAGCGGGGTATT |
| AACGCATATAATTGAGAATCGCCAGTAATTCTGTCAAAGCCA |
| CTCCGTGAAGTCTGAGAGACTACAAACTTTTTCAACAATAGT |
| GAATTCATACAAAGAAACCACCATTAATTACATTTTTGAGTA |
| AAATCACTGGCTGAGAGCCAGCAGTTGAAAGGAATATTAACA |
| TCAATGGATTGGAAATACCTACACCAACAGAGATAAGCCATT |
| CTGTGGGACGTAATCATGGTCATTTGATGGTGGTTGATCCCC |
| CTCGTATGTGGTAATGGGTAAAGAGCGGTGCCGGTAGCCGGG |
| GCGGTACGTTGCTGATTGCCGTTTAGAACGTCAGCTAGTGAT |
| CGACATACCCATCTGCCAGTTTGGAATTTGTGAGAGTGTAGA |
| CCCAAGCCACCCCGGTTGATAATTTTTGTTAAATCTAAAACT |
| ACGCAAGTTTTGCGGGAGAAGCCAAAGGGTGAGAAAAAAACA |
| TCAGAAGACGGTCTTTACCCTGATGGTCAATAACCCGAGAAT |
| ATACCAACAGATTCATCAGTTGACCCTCGTTTACCCGGAACA |
| CGGGAGAAGCCTTTATTGAGTCCCTCATAGTTAGCCCAAATATTTTGTT |
| TTAGTTACCAATCGAATAAACCAACTTTCAACAGTTTCAGCGATGCGTT |
| TCATTTGATGAAACCGTCGCTACTAAAGGAATTGCCGCAGAGCATTTCA |
| TCTGACCGACCAGTTAGCCCTTTATCAGCTTGCTTTCGAGGTAATGCGC |
| CGGCAAATTTGCCCCTATTAAACCGATAGTTGCGCTTCACCGAGAGAGT |
| TGGTCTGCCTCATAGCTGGAGGGCCGCTTTTGCGGGATCGTCACAACCA |
| TTTCTCCGTACAGCTTTGCCGCATCGGAACGAGGGAGCCGCCATAACCT |
| AGACAGTCTGAGTATGAGAGATACGTAATGCCACTACGAAGGAAGGTAG |
| CTATATTTTTGACCATCATACGCAAAAGAATACACTTCCATATTCCCAA |
| CGAGAAATCAACGTTGCGATTTCGCCTGATAAATTGTGTCGAAATGAAT |
| GCGTCCAAGCGACAGAATCAATCACCAATTGCCATGCAAGCCCAATAGG |
| AAACCAACCTTATCAACCAATCAATAATAATTTACCGTTTTT |
| TCAGCATCAGACGACGACAATGAATATATAATATCCAGACGTTAGTAAA |
| ACGCTCAACAAATTGCCTGTTTAGTATCTCATAATATTTAAC |
| GAATTTATAAGACGTTGAAAACATAGCGAATTTTCTTTTAAC |

| |
|---|
| ACATTATGTTATTACTCGTATTAAATCCTTACAAAAGGAGCG |
| TAGGAGTTGAGGAAGGTTATCATATCTGAGAAGTATCCAAAAAAAAGGC |
| CCGCCTGAGAGGTGACGAACCACCAGCACGCCATTAAATGAA |
| GGGTACCAGCCTCCCGCGTGCCTGTTCTCGGGGGTCTGTTTC |
| CAGTGTGGCCCCCTGCATCAGCATCAGATTTCACGATAACCGATATATT |
| TCACTGTGGGCGCGCAGGCGCTTTCGCAGCATCAGTTCTTTG |
| GAAGGGTAGCCGCACGACATAAAAAAATTTGGGCGGGCAAAC |
| TGGGCGCATGGGATGGGAACAAACGGCGACCCGTCGGGACGA |
| GCCATCGAGCTCATTTTTTAAATATTTTGTGAGCGTAAAGACTTTTTCA |
| AGCATGTATCGATGCTGAGAGTCTGGAGAGGCTATGAAAAGC |
| TTATGACAGCTAAATAAGCAATAAAGCCGCAAAGATATTTCA |
| GACCATATTTAAACTATTCATTGAATCCCTGCGGAATTATAG |
| GAGAGAAAGACGACGATAAAAGCATAGTGATAGCGTTATACCAAGCGCG |
| TAAGTAAGGTAAATTCACCGTCACCGAAAAGAAACGCAAAGAAAAGGTG |
| AGGGTGCTATCTTAGCCGAACAAAGTTAACTGAACACCCTACCATTAGA |
| TTTGAATGCGTTATACAGTAGGGCTTTTCATCTTCTGACCGTATATATT |
| TAAATAGCTTAGATTCAAAATCATAGTTACCTTTTTTAATGTAACAATT |
| CAGCACTAACAACTAATAGTGCCCGAACCATTTTGCGGAAAAATATCTT |
| AATATAGATAAAACCAACAGTGCCACAAAGCGTAAGAATACCGAACCCT |
| AGATACATAATATCGAAAAACGCTCACCCTTATAAATCAAAGCCGAAAT |
| TATCACTGCGCGCCTGTGCGCGTCCGTGGAGCTCGAATTCGATCCAGCG |
| GCAGCCAATCCGCCTGCCCTGCGGCTCAGCAGCAACCGCATTGTGGTGC |
| AGAGACGTAAAAAAAAAGTTAAACGAGGTGAAGGGATAGCTGGATAGAC |
| GTTCTAACAAGAGACAATCATATGTAAATCACCATCAATACCAGGCCGG |
| GCATCAGAGCATAACCTGTAATACTTCATTTGGGGCGCGAAATGTTTAG |
| CTGCTTTTGCAAAAGAAGTCTCAAATGCAATCAAAAATCACAAAATAGC |
| AGATGAAAATCTACTACAGGTAGAAACCAGAACGAGTAGTTTGCCCTGA |
| ATATCAGAGAGAGAATTCCAAGAGAACAAAGTCAG |
| GTGATAAATAAGCCCTGAGAAGATAAATTTAATGG |
| GTGAATAACCTTAAATTTTAAAAGGAAACAGTACA |
| ATTAGTCTTTAAGGATATTACCGCGTGGCACAGAC |
| GTTCCAGTTTGGCATCACAGTTGAAGAATAGCCCG |
| GCCATCCCACGCATCAGGCGGCCAGAATGCCAACG |
| GGATCAAACTTAGTAGGTCACGTTCTCACGGAAAA |
| AATGCCGGAGAGATTCGGTTGTATGATATTCAACC |
| AGTAGTAGCATTAAAGTTCAGAAGCTGAAAAGGTG |
| CTTTAATCATTGCGCTGACCAACAAATTGGGCTTG |
| TATGGTTGTATGTTCATGATTAAGACTCCTCAAAATCACCAGTAGCACC |
| CACAAGAACAGAGAGTCAAAAATGAAAATAAATAACGGAATA |
| AAATAAGCAAGACAGTAAATGCTGATGCAACAACATGTAATTTAGGCAG |
| TGTAAATAAACATCCCTGAGCAAAAGAAGAGCTTAGGTTGGG |
| GAACTGAAATAAAACAGATTCACCAGTCACTAAAGCATCACCTTGCTGA |

| |
|---|
| GAGTCCACAGCAGGGCAAGCGGTCCACGCTCGTCTGAAATGG |
| GCTTACGACGGAACCCTCCGGCCAGAGCACCATAAACATCCCTTACACT |
| CTGCTCAGCCATGTGGAAACAATCGGCGAACCGTTTTTTCGT |
| CTATTTTATGTGTAATATTTTAAATGCAATAAACAGGAAGATTGTATAA |
| CCAATAAATTAGATGCGAACGAGTAGATTTAGGATAAAAATT |
| TACCTTAAACAAAGGGATATTCATTACCCAACATTCAACTAATGCAGAT |
| AGGGATACTTTTCAGGCATCGGCATTTTCGGTCATAGCCCCAGAAATTA |
| AACCCATTTCCTTATTAGCGTTGAAACCTTAGCAAGGCCGGAAACAGAG |
| AACACTGAGTTTCGGCCAAAGGGTTATTTTGTCAC |
| CATTCCACAGACAGTAAGCCCTAAATTGAGCGCTA |
| GTCTTTCCCATCCTCGGTTTATCAACAATAGATAAGTCCTAACTTACCA |
| TGAATTTAAGAACAAGAAAAAAAGTACCTTCGAGCCAGTAATAACATCG |
| GGGATTTTGCTAAAACCGGAAATAATACCGACCGT |
| AATAGAAAGGAACAATTAATTATCAATATATGTGA |
| GAAAATCTTAGACTTTATTAGAGCCGTCAATAGATAATACGAAGGCGGT |
| TCCAAAACCATTTGAGGATTTGTCAGTTATATCAAACCCTCAATATCAT |
| TTAATTGTATCGGTAAAACATGATTTTGAATGGCT |
| TTAAACAGCTTGATAGAACGTATGGGTTGAGTGTT |
| CCCACGCGTCATACTCACTCTGTGGTGCTGCGGCCAGAATTGGTTGCGG |
| CGGTCGCATGCGGCGGGCCGTTGCCGGGTCAGCAAATCGTTAACTGAAA |
| TGCAGGGAGTTAAAGTGTCCACTACCGTCGGTGGT |
| GCAGCGAAAGACAGCCAGCAGCCCGCAGAAACAGC |
| TGAGGACAGTAACAGATGGCCTTCCTGTAGCCAGCTTTCAGCAACGGTA |
| TGAGGAATTTCAACATTAAATGTTAAAATTTAAATTGTAAACGTGTATC |
| TTAAACGGGTAAAATCTACAACAAGCTGATAAATT |
| CTAAAACGAAAGAGAGGCAAGTCAATTCTACTAAT |
| CCAGCGATCCAATACCTTTGCCAGAGGGGGTAATAGTAAAGTAAACGAA |
| AAACAAACAATGTTTAGACTGAAGAGCAGCCAAAAGGAATTACGAGCAA |
| GGAGATTTGTATCATTAAGAAGAGTTTAATTTCAA |
| GCCACCCCCTCAGACCAGAGCCACCACCTAATCAACAGAACC |
| GCAACAGCAGAACAGCGAAAAACCGTCTGGACTCCTTGACGC |
| ACATTATGTTAATACAGGACGTTGGGAACTGGCTCTTTAGGA |
| GTTAATGCAGAACGCGCCTGCTGTCTTTGTACCGCACTCAACAACATGT |
| ATCAAAAATAATTCGCGTCTTGACCGTAATCGTAACCGTGAATAGGAAC |
| GCAACATTTACGCATACCAGCTCACCAGTACAAACTACAACGGAATTCA |
| TGAATTAATTGACGACCGATTGAGGGAGACAAAAGAATTAGA |
| GCAGATACCGAAGCCAATGAAATAGCAAAATAATAAGGAAAC |
| TTAGACTGTAGCGCGTTTTAACCGCCTCTCAGAGCCACCATTGCCTTTA |
| AATCAATAGAAAAACCTTTTTAAGACACCACGGAA |

Handle staples for myosin and the actin binding domain of α actinin. Sequences in italic indicate the handle site. Position numbers in the list correspond to Fig. 1b.

| | |
|---|---|
| Handle for myosin at position 1 | GCTTTTGATGATTCCAGTAAGCGTCATA *TTCCTCTACCACCTACATCAC* |
| Handle for myosin at position 2 | ATTTTGCACCCATTAAATCAAGATTAGT *TTCCTCTACCACCTACATCAC* |
| Handle for myosin at position 3 | GTAGATTTTCAGAACAGAAATAAAGAAA *TTCCTCTACCACCTACATCAC* |
| Handle for myosin at position 4 | GTGCCAGCTGCATTCCAGTCGGGAAACC *TTCCTCTACCACCTACATCAC* |
| Handle for myosin at position 5 | CCAGCTGGCGAACGGGCCTCTTCGCTAT *TTCCTCTACCACCTACATCAC* |
| Handle for myosin at position 6 | TTTGATAAGAGGGAGTACCTTTAATTGC *TTCCTCTACCACCTACATCAC* |
| Handle for α actinin at position 1 | *CTCTCCTCTCCACCATATCCA* CATGGCTTTTGATGATTCCAGTAAGCGT |
| Handle for α actinin at position 2 | *CTCTCCTCTCCACCATATCCA* TGCTATTTTGCACCCATTAAATCAAGAT |
| Handle for α actinin at position 4 | *CTCTCCTCTCCACCATATCCA* TGTCGTGCCAGCTGCATTCCAGTCGGGA |
| Handle for α actinin at position 5 | *CTCTCCTCTCCACCATATCCA* TACGCCAGCTGGCGAACGGGCCTCTTCG |
| Handle for α actinin at position 6 | *CTCTCCTCTCCACCATATCCA* TCCTTTTGATAAGAGGGAGTACCTTTAA |

To label GNP to myosin at position 3, we used the handle below.

*AAAAAAAAAAAAAAAAAA* TTGCGTAGATTTTCAGAACAGAAATAAAGAAATTCCTC TACCACCTACATCAC*AAAAAAAAAAAAAAAAAA*
